# Production of Offspring from Azoospermic Mice with Meiotic Failure: Precise Biparental Meiosis within Halved Oocytes

**DOI:** 10.1101/2021.09.03.458818

**Authors:** Narumi Ogonuki, Hirohisa Kyogoku, Toshiaki Hino, Yuki Osawa, Yasuhiro Fujiwara, Kimiko Inoue, Tetsuo Kunieda, Seiya Mizuno, Hiroyuki Tateno, Fumihiro Sugiyama, Tomoya S. Kitajima, Atsuo Ogura

**Affiliations:** Bioresource Engineering Division, RIKEN BioResource Research Center, Ibaraki 305-0074, Japan; Laboratory for Chromosome Segregation, RIKEN Center for Biosystems Dynamics Research, Kobe, Hyogo 650-0047, Japan; Graduate School of Agricultural Science, Kobe University, Kobe, Hyogo 657-8501, Japan; Department of Biological Sciences, Asahikawa Medical University, Asahikawa, Hokkaido 078-8510, Japan; Graduate School of Comprehensive Human Sciences, University of Tsukuba, Tsukuba, Ibaraki 305-8575, Japan; Laboratory of Pathology and Development, Institute for Quantitative Biosciences, The University of Tokyo, Yayoi, Bunkyo-ku, Tokyo 113-8657, Japan; Graduate School of Life and Environmental Sciences, University of Tsukuba, Tsukuba, Ibaraki 305-8572, Japan; Faculty of Veterinary Medicine, Okayama University of Science, Imabari, Ehime 794-8555, Japan; Laboratory Animal Resource Center and Trans-border Medical Research Center, Faculty of Medicine, University of Tsukuba, Tsukuba, Ibaraki 305-8575, Japan; RIKEN Cluster for Pioneering Research, Wako, Saitama 351-0198, Japan

**Keywords:** azoospermia, fertilization, meiosis, oocyte, spermatocyte

## Abstract

While the large volume of mammalian oocytes is necessary for embryo development, it can lead to error-prone chromosomal segregation during meiosis. Conversely, we hypothesized that smaller oocytes would have a great unidentified potential to stabilize unstable meiosis and improve the development of the resultant embryos. Here, we show that reducing ooplasmic volume can rescue highly error-prone fertilization using primary spermatocytes by preventing segregation errors of chromosomes during biparental meiosis. High-resolution live-imaging analysis revealed that erroneous chromosome segregation occurred in most (90%) spermatocyte-injected oocytes of normal size, but could be ameliorated to 40% in halved oocytes. The birth rate improved remarkably from 1% to 19% (*P* < 0.0001). Importantly, this technique enabled the production of offspring from azoospermic mice with spermatocyte arrest caused by STX2 deficiency, an azoospermia factor also found in humans. Thus, contrary to popular opinion, oocytes inherently have a strong potential for precise meiotic divisions, which can be evoked by reduction of the ooplasmic volume. Their potential might help rescue cases of untreatable human azoospermia with spermatocyte arrest.

## INTRODUCTION

Fertilization is the process whereby female and male gametes (oocytes and spermatozoa) unite to form a zygote. From the standpoint of their genomes, the oocyte and spermatozoon are equivalent, but their history and cell type are quite different. Oocytes acquire their large cytoplasm (the ooplasm) during oogenesis to store all the components necessary for embryogenesis, including organelles, proteins, metabolites, mRNAs, and other molecules. By contrast, the contribution of spermatozoa to zygote formation and embryonic development is largely limited to deposition of the paternal genome and oocyte activation. Consequently, simple injection of a spermatozoon or even the sperm head (nucleus) into a mature oocyte results in normal fertilization, leading to embryo development and birth of offspring (Ogura et al., 2005; Palermo et al., 1992). Fertilization of oocytes does not even require mature sperm nuclei, because injection of nuclei from immature spermatozoa (spermatids) is sufficient for normal fertilization and embryo development to term (Ogura et al., 2005). Indeed, in one clinical study, 90 babies were born following round spermatid injection, without any significant adverse effects (Tanaka et al., 2018).

Therefore, a large ooplasm helps determine the embryo’s developmental potential. However, it is known that this feature does not always provide benefits for development. We and others have shown that a large ooplasm is linked to error-prone chromosomal segregation, by analyzing high-resolution images of meiotic chromosomes in oocytes with artificially increased or decreased ooplasmic volume (Kyogoku and Kitajima, 2017; Lane and Jones, 2017). Thus, the evolution of a particular ooplasmic mass in a species might have arisen as a delicate trade-off between meiotic fidelity and post-fertilization developmental competence. In our analysis, it was clear that a large ooplasm was detrimental, because the meiotic chromosomes showed frequent abnormal behavior, which could have led to aneuploidy and embryonic death. Conversely, one can postulate that a smaller ooplasm might be more beneficial than a larger one in terms of chromosomal behavior, but there is no evidence for this because intact oocytes undergo meiotic divisions normally during oogenesis and fertilization in experimentally tractable animal models such as mice.

As mentioned above, normal diploid embryos can be obtained using spermatids because they are already haploid, as are mature spermatozoa. However, the use of primary spermatocytes for fertilization is considered to be ineffective because they are premeiotic germ cells. Theoretically, the chromosomes of primary spermatocytes might be able to contribute to the construction of diploid embryos after two meiotic divisions within oocytes. Indeed, we and another group have reported the birth of mice following spermatocyte injection into oocytes, but the success rates were low at 1% to 3% per embryo transferred (Kimura et al., 1998; Ogura et al., 1998). This was mostly caused by the death of embryos shortly after implantation. When we observed the reconstructed oocytes at metaphase II (MII), there was a high incidence of chromosomal aberrations (Miki et al., 2006; Ogura et al., 1998). Since then, there have been no technical improvements in spermatocyte injection. However, the use of primary spermatocytes for conception should be explored, given that many cases of nonobstructive azoospermia in humans are associated with spermatogenic arrest at the primary spermatocyte stage (Enguita-Marruedo et al., 2019).

Based on these findings, we expected that reducing the ooplasmic volume might improve the chromosomal integrity of spermatocyte-injected oocytes and increase the survival rate of the resultant embryos. In this study, by employing high-resolution live-imaging techniques, we analyzed the segregation patterns of the maternal and paternal chromosomes within spermatocyte-injected oocytes with or without reduction of the ooplasm. Furthermore, we examined whether such reduction could improve the birth rate following spermatocyte injection and whether this technology could be applied to azoospermic mice having a mutation causing spermatocyte arrest.

## RESULTS

### Reduction of the Recipient Ooplasmic Volume Increases the Rate of Normal Diploidy in Spermatocyte-injected Oocytes

Fertilization with primary spermatocytes was achieved by injecting a spermatocyte nucleus into immature oocytes at the germinal vesicle (GV) stage followed by arrest at the metaphase of meiosis I (MI) induced by cytochalasin D treatment (**Figure 1A**). Here, the maternal and paternal (spermatocyte-derived) chromosomes were synchronized, forming a single chromosomal mass. After removal of cytochalasin D, they underwent meiotic division with protrusion of the first polar body to reach the MII stage (**Figure 1A**). This reconstructed MII “zygote” could be activated artificially to resume meiosis and form a one-cell embryo having one zygotic nucleus (**Figure 1A**). To test the developmental ability of reconstructed embryos, we transferred these MII chromosomes to freshly prepared enucleated MII oocytes (Ogura et al., 1998).

**Figure 1.**
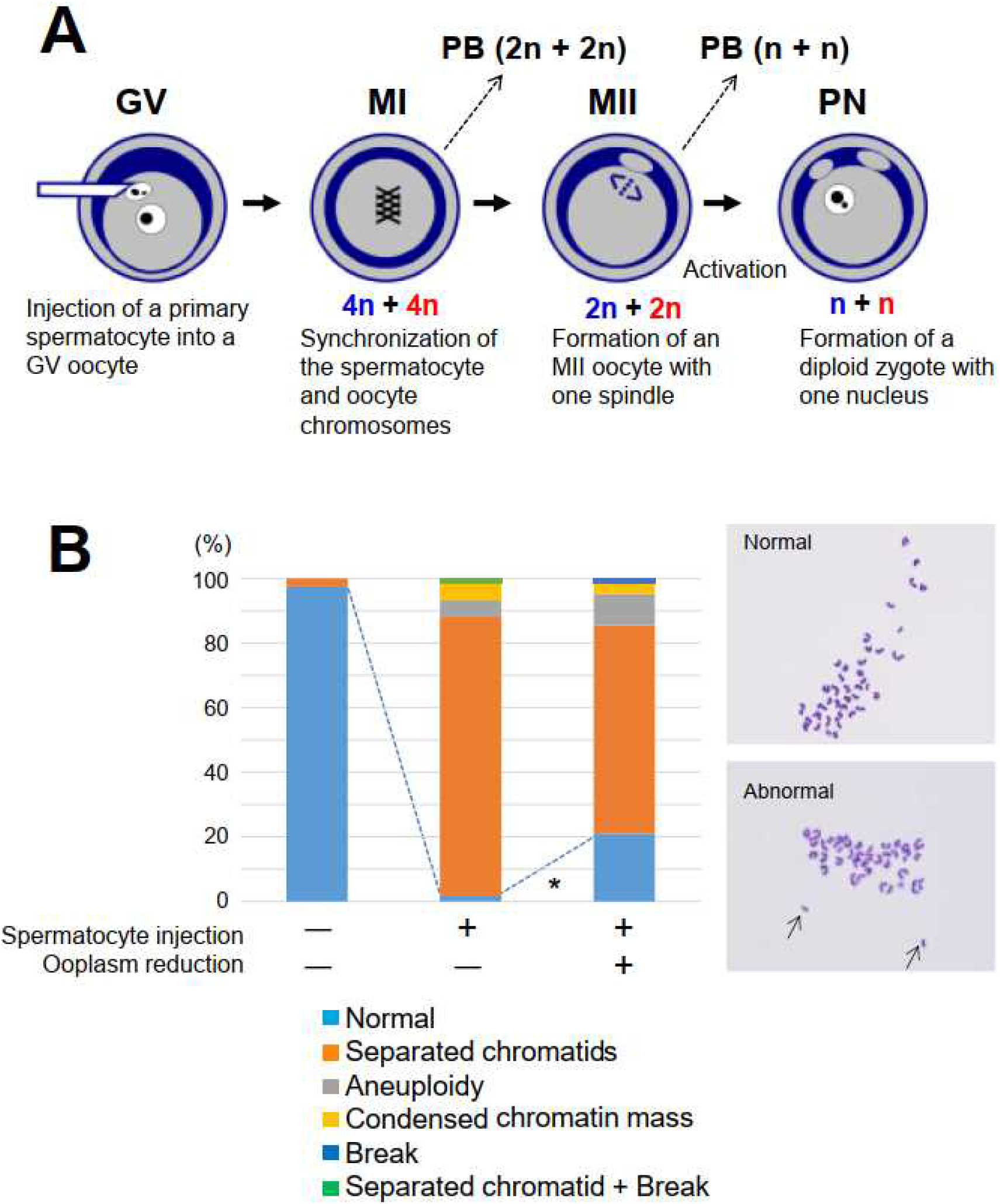
Fertilization with Primary Spermatocytes. (A) The scheme of construction of a diploid fertilized oocyte using a primary spermatocyte and a GV-stage oocyte. The chromosomes of the spermatocyte and the oocyte are intermingled at MI to form a single chromosomal mass. (B) Chromosomal analysis of MII oocytes that had been injected with primary spermatocytes. In the spermatocyte-injected groups, normality was improved by reducing the ooplasm mass (\**P* < 0.005 by Fisher’s exact probability test). Arrows in the right figure indicate prematurely separated chromatids. For the exact numbers in each case, see also **Table S1**. PB, polar body; GV, germinal vesicle; MI, meiosis I; PN, pronuclear stage.

Recipient oocytes with half the normal ooplasmic volume were prepared by aspiration using a large glass pipette (**Movie S1**). Using these halved oocytes, we first analyzed the chromosomal integrity of spermatocyte-injected oocytes. In control oocytes without spermatocyte injection (i.e., oocyte chromosomes only), the proportion of oocytes with normal chromosomes was 97% at MII (**Figure 1B and Table S1**). In spermatocyte-injected oocytes with intact ooplasm, the proportion of normal chromosomes was decreased significantly to 2% (1/59, *P* < 0.0001) (**Figure 1B and Table S1**). The most frequent abnormality (86%, 51/58) was the presence of prematurely separated sister chromatids (**Figure 1B and Table S1**). When spermatocytes were injected into half-sized oocytes, the proportion of MII oocytes with normal chromosomes improved significantly to 13/62 (21%; *P* < 0.005, vs the intact cytoplasm group) because of the decreased number of separated chromatids (**Figure 1B and Table S1**). Thus, while chromosomal normality was largely lost during meiosis I in spermatocyte-injected oocytes, chromosomal aberrations could be prevented in a significant proportion of oocytes by reduction of the ooplasmic mass.

### Reduction of the Recipient Ooplasm Corrects the Behavior of Spermatocyte-derived Chromosomes During Meiosis

Next, we sought to study how chromosomal behavior was influenced by the ooplasmic volume and which of the two parental (maternal or paternal) chromosomes was more vulnerable to the stress of biparental meiosis. The high-resolution three-dimensional (3D) live imaging system reported in our previous studies was employed for analyzing the chromosomal behavior during meiosis I (Kitajima et al., 2011; Kyogoku and Kitajima, 2017). To this end, it was essential to discriminate the origins of the chromosomes via fluorescence microscopy. Interestingly, the paternal (spermatocyte-derived) chromosomes could be distinguished from the maternal chromosomes by the relatively lower fluorescent intensities of the histone H2B-mCherry marker (**Figure 2A**). Our 3D visualization of individual chromosomal positions showed that biparental meiosis exhibited more frequent misalignment of paternal chromosomes at late MI, compared with the maternal chromosomes (**Figures 2B, C and Movie S2**). Halving ooplasmic volume significantly reduced the number of misaligned paternal chromosomes (**Figures 2B, C and Movie S3**), an effect that we expected based on our previous observations (Kyogoku and Kitajima, 2017). Thus, paternal chromosomes are susceptible to errors in ooplasm-hosted biparental meiosis, which can be tuned by reducing the ooplasmic volume.

**Figure 2.**
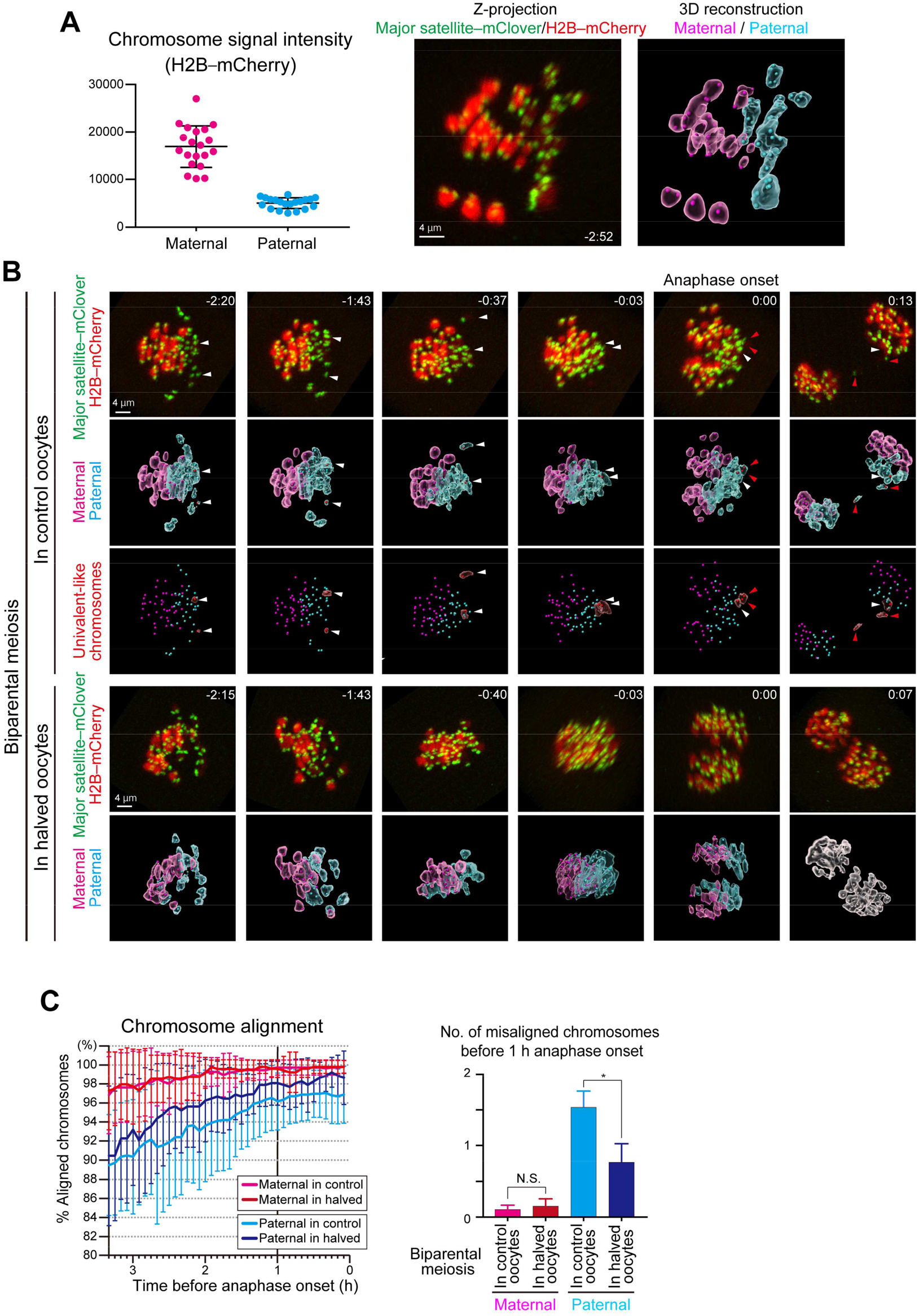
Live Imaging of Biparental Meiosis. (A) Identification of parental origin of the chromosomes was distinguishable based on H2B-mCherry fluorescent intensities (paternal chromosomes exhibit lower intensities). The z-projection image shows major satellite–mClover (centromeres, green) and H2B-mCherry (chromosomes, red). Time from anaphase onset is shown in h:min. Scale bar = 4 µm. The 3D-reconstructed image shows maternal (magenta) and paternal (cyan) chromosomes. Spots indicate centromeres. (B) Chromosome tracking in 3D. The reconstructed images are viewed from the side of the metaphase plate. Signals are interpolated in the Z axis for visualization. White and red arrowheads, as well as red surfaces, indicate univalent-like chromosomes that underwent unbalanced predivision (premature segregation of sister chromatids). Scale bar = 4 µm. (C) Halving the recipient ooplasmic mass rescued chromosome alignment. The numbers of misaligned chromosomes and their parental origin were determined in 3D (*n* = 39 and 17 oocytes). Error bars show the standard deviation. Student’s *t* test was used to compare means. \**P* < 0.05. N.S., not significant.

We then analyzed how biparental meiosis in normal-sized ooplasms results in chromosomal abnormality. Our technique of complete centromere tracking using 3D imaging (Kitajima et al., 2011; Sakakibara et al., 2015) enabled us to demonstrate that 89% of biparental meiotic divisions showed errors in chromosomal segregation at anaphase I (**Figures 2B, 3A and Movie S2**). Almost all of the errors were of spermatocyte origin (**Figure 3A**). Categorization of anaphase trajectories showed that predominant error patterns were balanced and unbalanced predivisions (premature segregation of sister chromatids at MI) (**Figure 3B**), consistent with our observation of separated chromatids in MII spreads (**Figures 1B and Table S1**).

**Figure 3.**
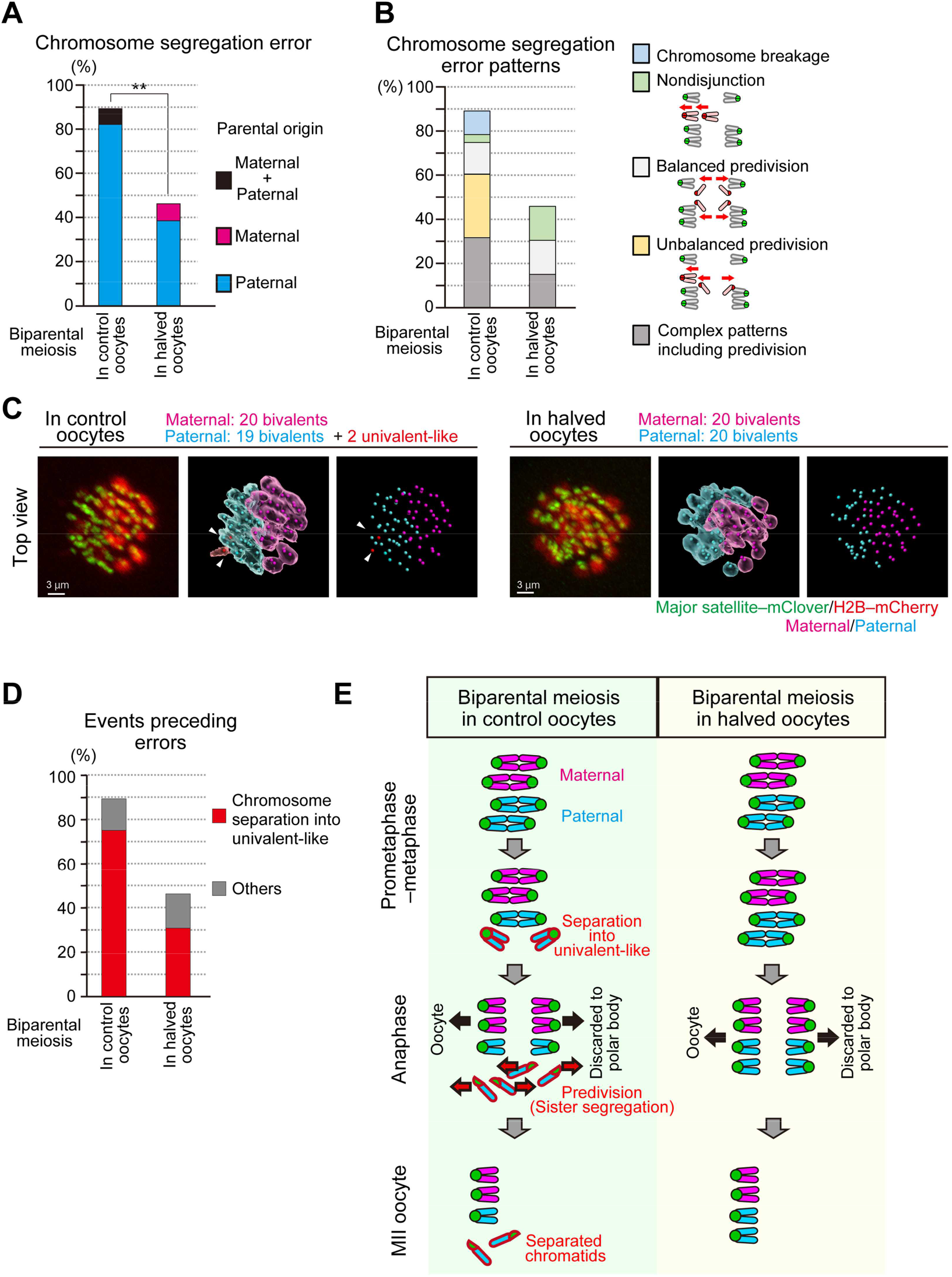
Halving the Recipient Ooplasm Prevents Premature Separation of Paternal Chromosomes in Biparental Meiosis. (A) Halving the recipient ooplasm mass reduced chromosome segregation errors. Errors were determined by tracking all chromosomes at anaphase (*n* = 39 and 17 oocytes) (See also **Figure 2B**). The parental origin of errors is shown. Ooplasmic halving significantly reduced the rate of errors (\*\**P* < 0.01). (B) Predivision was predominant in biparental meiosis. Chromosome segregation error patterns were categorized based on anaphase trajectories: nondisjunction (0:4 segregation), balanced predivision (2:2 sister chromatid segregation), unbalanced predivision (1:3 segregation including sister chromatid segregation), and complex patterns including predivision (multiple errors including sister chromatid segregation). Chromosome breakages (chromosomes lacking centromeres) were also observed. (C) Univalent-like chromosomes. Images were 3D-reconstructed as in Fig. 2B and viewed from the top of the metaphase plate. Red surfaces with white arrowheads indicate univalent-like chromosomes. Scale bar = 3 µm. (D) Halving the ooplasm volume suppressed the premature separation of paternal chromosomes. Oocytes were categorized based on whether the chromosomes exhibited premature separation into univalent-like structures prior to segregation errors. (E) Summary of biparental meiosis. In normal-sized oocytes this frequently exhibits premature separation of paternal chromosomes into univalent-like structures. These chromosomes undergo predivision (premature segregation of sister chromatids), and thus result in separated chromatids in MII oocytes. Halving the ooplasmic volume reduced such errors.

Predivisions are error patterns observed following premature separation of bivalent chromosomes into univalents during the prometaphase and metaphase in naturally aged oocytes (Sakakibara et al., 2015). Therefore, we carefully analyzed the prometaphase–metaphase trajectories of the chromosomes that underwent segregation errors in biparental meiosis. This analysis revealed that most of the errors were preceded by premature separation of bivalent chromosomes into univalent-like structures (**Figures 2B, 3C and Movies S2, S3**). Importantly, decreasing (halving) the recipient ooplasm mass significantly suppressed the premature bivalent separation of chromosomes (75% in controls vs 31% in halved oocytes) (**Figure 3D**) and chromosome segregation errors (89% in control vs 46% in halved oocytes) (**Figures 3A, B, and D**). Thus, the chromosomal aberrations found in spermatocyte-injected oocytes were largely attributable to the premature separation of spermatocyte-derived chromosomes, and about half of such aberrations could be prevented by reducing the size of the recipient ooplasm (**Figure 3E**).

### Reduction in the Recipient Ooplasm Improved the Birth Rates Following Spermatocyte Injection

Next, we examined whether reduction of the ooplasm volume could improve the developmental ability of spermatocyte-derived embryos. When we reconstructed embryos using normal-sized oocytes and transferred them into recipient females, only 1% (1/96) developed into offspring (**Figure 4A and Table S2**), consistent with our previous reports (Miki et al., 2006; Ogura et al., 1998). By contrast, when we used halved oocytes, 19% (17/90) of the reconstructed embryos developed into live offspring, achieving a nearly 20-fold improvement (*P* < 0.0001, **Figure 4A and Table S2**). The pups born by this improved method had body and placental weights within the normal ranges (**Figure S1**). We allowed three male pups to grow into adults and confirmed that they were all fertile by mating them with normal female mice.

**Figure 4.**
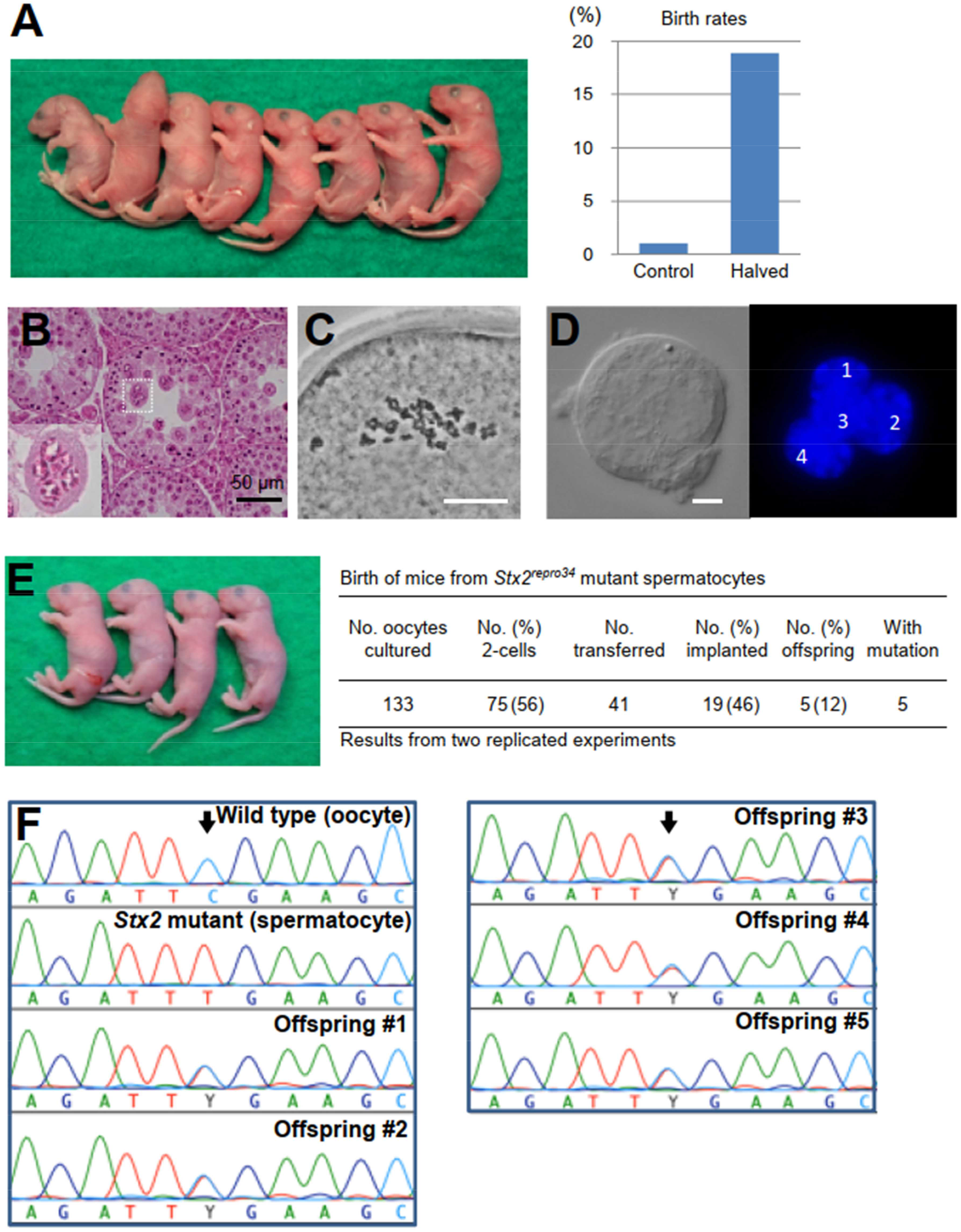
Birth of Spermatocyte-derived Offspring Following Embryo Transfer. (A) Mouse pups born following spermatocyte injection (left) and the birth rates following embryo transfer (right). For detailed results, see also **Table S2**. (B) Histology of the testis of a *Stx2*^*repro34*^ mouse. Arrowheads indicate multinucleated cells containing spermatocyte-like nuclei. There are no spermatids or spermatozoa. Bar = 50 µm. (C) An MII oocyte injected with a putative spermatocyte nucleus from a multinucleated cell in a *Stx2*^*repro34*^ mouse testis, showing the typical paired meiotic chromosomes. Bar = 20 µm. (D) A multinucleated cell isolated from a *Stx2*^*repro34*^ mouse testis, showing four nuclei. Differential interference contrast (left) and Hoechst-stained (right) images. Bar = 10 µm. (E) (left) mouse pups born following microinjection with putative primary spermatocyte nuclei isolated from multinucleated cells; (right) birth rate of pups following *Stx2*^*repro34*^ spermatocyte microinjection. (F) Genomic sequencing confirming the origin of pups from *Stx2*^*repro34*^ spermatocytes. Arrows indicate the expected point mutation of *Stx2*^*repro34*^. Y indicates a hybrid status with T and C bases.

### Spermatocyte Injection Rescued Azoospermia Caused by Meiotic Arrest

Finally, we applied this improved spermatocyte injection method to mouse strains with azoospermia caused by spermatogenic arrest at the primary spermatocyte stage. If the chromosomes of spermatocytes are functionally and structurally intact, we surmised that their normal meiotic divisions might be induced by the meiotic machinery of recipient oocytes. We performed our studies on *Stx2*^*repro34*^ mice (hereafter, *repro34* mice) that carry a mutation in the *Stx2* (syntaxin 2) gene induced by *N*-ethyl-*N*-nitrosourea (ENU) mutagenesis (Fujiwara et al., 2013). Its human homologue, *STX2*, has been identified as a causal factor of nonobstructive azoospermia (Nakamura et al., 2018). Both mouse *Stx2* and human *STX2* mutations are characterized by the formation of large syncytial spermatocytes because of their inability to maintain intercellular bridges (Fujiwara et al., 2013; Nakamura et al., 2018) (**Figure 4B**). We confirmed that the nuclei within these syncytial cells of *repro34* mice were derived from spermatocytes by examining their prophase I chromosomes following injection into MII oocytes (**Figure 4C**). We reconstructed embryos using the nuclei collected from these syncytial spermatocytes (**Figure 4D and Movie S4**). After 41 embryos were transferred into recipient females, five pups (four female and one male) were born (**Figure 4E**). All of these pups carried the point mutation in the *Stx2* gene (**Figure 4F**). We also applied this technique to spermatocytes from *Exoc1* (exocyst complex component 1)-deficient mice that also show syncytial spermatocytes (Osawa et al., 2021). Three pups (two female and one male) carrying the mutation were born at term (**Figure S2**). All the eight pups derived from *Stx2*- or *Exoc1*-deficient spermatocytes looked normal in appearance and their body and placental weights were within normal ranges, except for the body weight of pups from *Stx2*-deficient spermatocytes (**Figure S1**). They grew into normal-looking adults and were proven to be fertile.

### Chromosomal Analysis of Offspring Born Following Spermatocyte Injection

As described above, all the pups born following the injection of wild-type spermatocytes or mutant spermatocytes grew into fertile adults. We then analyzed their chromosomal constitution in detail by multicolor fluorescence *in situ* hybridization (FISH). Among the three male mice derived from wild-type spermatocytes, two had a normal karyotype, but one had XYY sex chromosomes (**Figure S3**). Among the five mice (four female and one male) derived from *Stx2*-deficient spermatocytes, two female mice and one male mouse were normal, but one female had an XO chromosome and another had a shortened X chromosome (**Figure S4**). Among the three female mice derived from *Exoc1*-deficient spermatocytes, one had an XO chromosomal configuration. No abnormalities were found in the autosomes of the mice examined.

## DISCUSSION

Here, we addressed whether reducing the recipient ooplasm could ameliorate the embryonic death rate caused by the meiotic errors that can occur in mammalian oocytes. To this end, we employed an assisted fertilization system using primary spermatocytes, which need simultaneous biparental meiosis within oocytes: namely, meiosis with doubled chromosomes. Following spermatocyte injection into halved oocytes, the proportion of normal chromosomes at MII increased from 2% to 21% and the birth rate increased from 1% to 19%. These results demonstrate unequivocally that reducing the mass of the ooplasm indeed helps to normalize chromosomal behavior, leading to better survival of embryos to term. It would be interesting to test whether this strategy could also correct the meiotic errors that are frequently found in oocytes from aged female mammals (Mihajlović and FitzHarris, 2018). In humans, these meiotic errors in oocytes are known to increase with advanced age and to reduce conception rates significantly (El Yakoubi and Wassmann 2017; Mikwar et al., 2020). In these errors, diverse mechanistic defects are involved, such as defects in chromosomal cross-over formation, cohesin loss and spindle deformation (Ma et al., 2020; Mihajlović and FitzHarris, 2018). We suspect that reducing the mass of the ooplasm might help rescue or prevent at least some of these defects.

The nearly 20-fold improvement in the birth rate following spermatocyte injection into halved oocytes was much better than we expected. It is known that meiosis in female and male mammals differs largely with respect to the underlying molecular mechanisms and cell cycle progression patterns, such as absence of the interphase between two meiotic divisions in female germ cells. Consistent with this, many strains of gene knockout mice carrying mutations of meiosis-related factors show male or female infertility (Biswas et al., 2021; Jamsai and O’Bryan, 2011). Our findings imply that the meiotic chromosomes of female and male germ cells have structural commonalities that allow mechanistic interchangeability between them. Nevertheless, most chromosomal aberrations were identified as of spermatocyte origin with a high incidence of premature sister chromatid segregation during meiosis I. Most of these errors were preceded by premature separation of bivalent chromosomes into univalent-like structures. Our results reveal a novel effect of ooplasmic reduction that can suppress premature separation of chromosomes, at least in the context of biparental meiosis. These suggest that spermatocyte-derived chromosomes are more vulnerable to physical or biochemical properties associated with a large ooplasm, such as spindle size, and ooplasmic dilution of nuclear factors (Kyogoku and Kitajima, 2017). The delayed alignment of spermatocyte-derived chromosomes to the MI spindle (**Figures 2B and C**) might also reflect these differences. In other words, maternal meiotic chromosomes might have evolved special mechanisms that efficiently avoid segregation errors in a large ooplasm.

Chromosomal analysis by multicolor FISH revealed that four of the 11 spermatocyte-derived offspring carried chromosomal abnormalities that were restricted to the sex chromosomes. There were no abnormalities in autosomes in any of the mice analyzed. This sex-chromosome-biased chromosomal aberration may be explained by the high embryonic lethality of autosomal aneuploidy, which might have selected embryos with normal autosomes for survival to term. It might also have resulted from yet undiscovered special characteristics of sex chromosomes, especially those derived from spermatocytes. All the abnormal patterns found in the sex chromosomes—XO, XX with a shortened X, and XYY—can be explained by segregation errors of spermatocyte XY chromosomes during meiosis I (see **Figures 3B and E**). It is known that sex chromosomes are prepared to undergo meiosis later than autosomes as they require the formation of the XY body (Kauppi et al., 2011). Therefore, it is possible that spermatocytes that had not completed this stage might have been selected for injection accidentally.

This study has practical implications for treating spermatogenic arrest caused by meiotic arrest. Given the complexity of meiosis, many genes are involved in its regulation, as revealed by mouse gene knockout models (Jamsai and O’Bryan, 2011). Defects in some of these genes might cause failure of meiosis and spermatogenic arrest at the primary spermatocyte stage. The results of this study will help identify the types of meiotic arrest that can be rescued or prevented by the spermatocyte injection technique we developed here. Such information would provide invaluable clues for human clinical research aiming to develop treatments for meiosis-related male infertility. In addition, we propose another important implication of this study: at present, complete *in vitro* gametogenesis is possible for female germ cells (Hikabe et al., 2016), but not for male germ cells. One of the major causes of this sex-specific difference in *in vitro* gametogenesis is the inability of male germ cells to undergo meiosis *in vitro*. If male primordial germ cell-like cells derived from induced pluripotent stem cells (Hayashi et al., 2011) could be cultured to form pachytene spermatocytes, injecting them into immature oocytes as substitute gametes might produce offspring by skipping *in vivo* male gametogenesis completely. This could be the ultimate strategy to enable conception in cases of male patients with germ cell loss. These scenarios could open up new methods for treating human male infertility, although there are a number of ethical and technical issues—for example, the high incidence of sex chromosome abnormalities—that need to be resolved before these strategies could be used by clinics offering assisted reproductive technology.

## MATERIALS AND METHODS

### Mice

All animal experiments were approved by and performed according to the principles of the Institutional Animal Care and Use Committees at RIKEN, Tsukuba and Kobe Branches. B6D2F1 and C57BL/6NCrSlc mice were purchased from Japan SLC. ICR mice were purchased from CLEA, Japan. *Repro 34* mice carrying the *Stx*^*repro34*^ mutation were introduced from Okayama University. The *repro34* mutation was induced in a C57BL/6J male and subsequently a congenic line with the C3HeB/FeJ background was created (Akiyama et al., 2008). Germline-specific *Exoc1* conditional knockout mice were generated by breeding *Exoc1*^*tm1c(EUCOMM)Hmgu*^ mice (Skarnes et al., 2011) with *Nanos3*-Cre driver mice (kindly gifted by Dr Y. Saga, RIKEN BRC RBRC02568), which express Cre in spermatogonia (Osawa et al., 2021; Suzuki et al., 2008).

### Collection of oocytes

Female B6D2F1 mice (9–12 weeks old) were injected with 7.5 IU of equine chorionic gonadotropin (eCG, ASKA Pharmaceutical, Tokyo,Japan). Forty-four to forty-eight hours after injection, fully grown oocytes at the GV stage were collected from large antral follicles and released into M2 medium supplemented with 150 μg/ml dibutyryl cyclic (dbc) AMP (Merck KGaA). After being freed from cumulus cells by pipetting, oocytes were cultured for at least 1 hours in MEM) Merck KGaA) supplemented with 50 μg/mL gentamicin, 0.22 mM Na-pyruvate, 1 μg/ml epidermal growth factor (EGF), 150 μg/ml dbc AMP, and 4 mg/ml bovine serum albumin (BSA), (mMEM) (Fulka and Langerova, 2014) at 37°C in an atmosphere of 5% CO_2_ in humidified air, until micromanipulation.

### Collection of primary spermatocytes

Spermatogenic cells were collected from the testes of male B6D2F1, C57BL/6N, and ICR mice (12–16 weeks old) by a mechanical method as reported in a previous study (Ogura and Yanagimachi, 1993). Briefly, the testes were placed in erythrocyte-lysing buffer (155 mM NH_4_Cl, 10 mM KHCO_3_, 2 mM EDTA; pH 7.2). After the tunica albuginea had been removed, the testes were transferred into a cold (4°C) Dulbecco’s phosphate-buffered saline (PBS) supplemented with 5.6 mM glucose, 5.4 mM sodium lactate, and 3 mg/ml BSA (GL-PBS) (Ogura et al., 1996). The seminiferous tubules were cut into small pieces using a pair of fine scissors and pipetted gently to allow spermatogenic cells to be released into the medium. The cell suspension was filtered through a 38-μm nylon mesh and washed twice by centrifugation (200*g* for 4 min). After gentle washing, the cells were resuspended in GL-PBS and stored at 4°C until microinjection.

### Micromanipulation

To make half-sized oocytes, oocytes at the GV stage were transferred to M2 medium (Merck Millipore) containing 7.5 μg/ml cytochalasin D (Merck KGaA) and 60 mM NaCl for 10 min at 37°C. All manipulations were performed under an inverted microscope with a Piezo-driven micromanipulator (PrimeTech). The zona pellucida was opened by piezo drilling and one-third to half of the ooplasmic volume was aspirated with an injection pipette (inner diameter 25 μm) at 37°C (**Movie S1**). After manipulation, oocytes were cultured in mMEM containing 7.5 μg/ml cytochalasin D and 40 mM NaCl at 37°C in an atmosphere of 5% CO_2_ in air. About 1–1.5 hours later, primary spermatocytes (pachytene to diplotene stages) were injected into oocytes that were induced to arrest at the MI stage by cytochalasin D. Oocytes were cultured in mMEM containing 7.5 μg/ml cytochalasin D and 40 mM NaCl for 2 hours at 37°C in an atmosphere of 5% CO_2_ in humidified air. After washing in mMEM, the oocytes were cultured for 14– 17 hours until they reached the MII stage. The karyoplasts containing chromosomes were removed and were then fused with fresh enucleated oocytes using Sendai virus (HVJ; Ishihara Sangyo Co., Ltd.) in Hepes-buffered CZB medium (Chatot et al., 1990) containing 7.5 μg/mL cytochalasin B. After manipulation, the oocytes were cultured in CZB medium containing 7.5 μg/mL cytochalasin B for 1 hours at 37°C in an atmosphere of 5% CO_2_ in humidified air until complete fusion occurred. Reconstructed oocytes were activated by culturing them in Ca^2+^-free CZB medium containing 8 mM SrCl_2_ for 20 min. After washing, the oocytes were cultured in CZB medium for 24 hours under 5% CO_2_ in humidified air at 37°C.

### Embryo Transfer

Embryos that reached the 2-cell stage by 24 hours were transferred into the oviducts of Day 1 pseudopregnant ICR strain female mice (9–12 weeks old). On day 19.5, recipient females were euthanized and their uteri were examined for live fetuses. In some experiments, live fetuses were nursed by lactating foster ICR strain mothers. After weaning, they were checked for fertility by mating with ICR mice of the opposite sex.

### Chromosome preparation of oocytes

The MII oocytes were treated with 0.5% actinase E (Kaken Pharmaceutical Co.) for 5 min at room temperature to loosen the zona pellucida and then treated with a hypotonic solution (1:1 mixture of 1.2% sodium citrate and 60% fetal bovine serum, FBS; Merck KGaA) for 10 min at room temperature. Chromosome slides were prepared using a gradual-fixation/air-drying method (Mikamo and Kamiguchi, 1983). Briefly, oocytes were treated with Fixative I (methanol:acetic acid:distilled water = 5:1:4) for 6–8 min and put onto a glass slide with a small amount of Fixative I. Then, the oocytes were treated with Fixative II (methanol:acetic acid = 3:1) for 2 min, followed by treatment with Fixative III (methanol:acetic acid:distilled water = 3:3:1) for 1 min. The slides were air-dried under conditions of 50%–60% humidity at 22–24°C. For conventional chromosome analysis, the slides were stained with 2% Giemsa (Merck KGaA) for 8 min. C-band staining was used to distinguish between structural chromosome aberrations and aneuploidy (Tateno and Kamiguchi, 2007).

### Chromosome analysis by multicolor fluorescence *in situ* hybridization (FISH)

Spleens were removed under sterile conditions from mice produced by spermatocyte injection. Lymphocytes were isolated from the spleen and incubated in a tissue culture tube at a cell concentration of 1 × 10^6^/ml in RPMI1640 (Nacalai Tesque) containing lipopolysaccharide (10 μg/ml, Merck KGaA), concanavalin A (3 μg/ml, Nacalai Tesque), 2-mercaptoethanol (50 μM, Nacalai Tesque), and FBS (6%) at 37°C under 5% CO_2_ in humidified air for 48 hours. Colcemid (KaryoMAX, Gibco) at a concentration of 0.02 μg/ml was added to the cell suspension for the last 2 hours of culture to arrest the cell cycle at metaphase. The cells were centrifuged at 420*g* for 5 min and resuspended in 3 ml of a hypotonic solution (0.075 M KCl). Twenty minutes later, 2 ml of Carnoy’s fixative (methanol:acetic acid = 3:1) was added to the cell suspension. Cells were centrifuged at 420*g* for 5 min and resuspended in 5 ml of fresh Carnoy’s fixative. Centrifugations and fixations were repeated three times. Chromosome preparations were made using a Hanabi metaphase spreader (ADSTEC). For multicolor FISH analysis, the chromosome slides were hybridized with 21XMouse (MetaSystems) according to the manufacturer’s protocol. For denaturation of chromosomal DNA, the slides were incubated in 2 × saline sodium buffer (SSC) at 70°C for 30 min and then treated with 0.07 M NaOH at room temperature for 1 min. The denatured slides were washed in 0.1 × SSC and 2 × SSC at 4 °C for 1 min each and dehydrated with a series of 70%, 95%, and 100% ethanol. Multicolor FISH probes were denatured at 75°C for 5 min and applied to the chromosome slides. After hybridization at 37°C for 48 hours in a humidified chamber, the chromosome slides were treated with 0.4 SSC at 72°C for 2 min, washed in 2 SSC with 0.05% Tween20 (Merck KGaA) at room temperature for 30 seconds, and rinsed in distilled water. For counterstaining, the slides were covered by a coverslip with DAPI/Antifade (MetaSystems). The chromosome slides were observed using fluorescent microscopy. Fluorescence images were captured using a high-sensitive digital camera (α7s, SONY). The images were imported into the ChromaWizard software (Auer et al., 2018) to assign fluorescence colors to each chromosome. Based on these fluorescence colors, the chromosome numbering was determined. Ten metaphase cells per mouse were analyzed for karyotyping.

### Chromosome analysis by Giemsa banding (G-banding)

When multicolor FISH analysis revealed a possible chromosome deletion, additional G-band staining was performed to identify the lost part of the chromosomes. The chromosome slides were treated with 0.025% trypsin (FUJIFILM Wako Pure Chemical Corporation) for 2 min at room temperature, washed in PBS, and stained with 4% Giemsa for 8 min. Deletion sites were determined according to the band pattern nomenclature of mouse chromosomes (Nesbitt and Francke, 1973).

### Live cell imaging

After linearization of the template plasmids, mRNA was synthesized using the mMESSAGE mMACHINE KIT (Ambion). The synthesized RNAs were stored at −80 °C until use. The *in vitro*-transcribed mRNAs (1.2 pl of 650 ng/ l major satellite-mClover (Miyanari et al., 2013) and 0.6 pl of 350 ng/ l H2B-mCherry) were microinjected into oocytes. These were cultured for 1 hours and then subjected to micromanipulation. Live cell imaging was performed as described (Kitajima et al., 2011; Sakakibara et al., 2015), with some modifications. Briefly, a Zeiss LSM710 or LSM880 confocal microscope equipped with a 40 × C-Apochromat 1.2NA water immersion objective lens (Carl Zeiss) was controlled by a multi-position autofocus macro (Politi et al., 2018). For centromere tracking (**Figure 3**), 19 confocal z-sections (every 1.5 μm) of 512 × 512 pixel x/y images covering a total volume of 35.4 × 35.4 × 28.5 μm were acquired at 200-second intervals for at least 10 hours after spermatocyte injection into oocytes expressing major satellite-mClover and H2B-mCherry. Centromere tracking was performed as described (Kitajima et al., 2011; Sakakibara et al., 2015). The parental origin of chromosomes was identified by the intensities of the chromosomes and the centromeres (lower fluorescent intensity for the spermatocyte chromosomes) (**Figure S2A**).

### Statistical analysis

The rates of chromosomal abnormalities and embryo development were evaluated using Fisher’s exact probability test. The body and placental weights of pups were evaluated using Student’s *t*-test.

## SUPPLEMENTAL INFORMATION

Supplemental information can be found at..

## ACKNOWLEDGMENTS

This study was supported by KAKENHI Grant Numbers 23500507 (N.O.), JP19H05758 (A.O. and T.H.), 20H05376 (H.K.), and 18H05549 (T.S.K.).

## AUTHOR CONTRIBUTIONS

N.O. and A.O. conceived the project. The project was developed jointly by A.O., T.K., S.M., H.T., T.S.K., and F.S. Experiments were carried out by N.O., T.H., H.K., Y.O., Y.F., and K.I. The paper was written by N.O., T.H., H.K., T.S.K., and A.O.

