## Supplementary material for "Production of Offspring from Azoospermic Mice with Meiotic Failure: Precise Biparental Meiosis within Halved Oocytes": Suppl figures

**This PDF file includes:**

Figures S1 to S4
Tables S1 to S2
Legends for Movies S1 to S4

**Other supplementary materials for this manuscript include the following:**

Movies S1 to S4

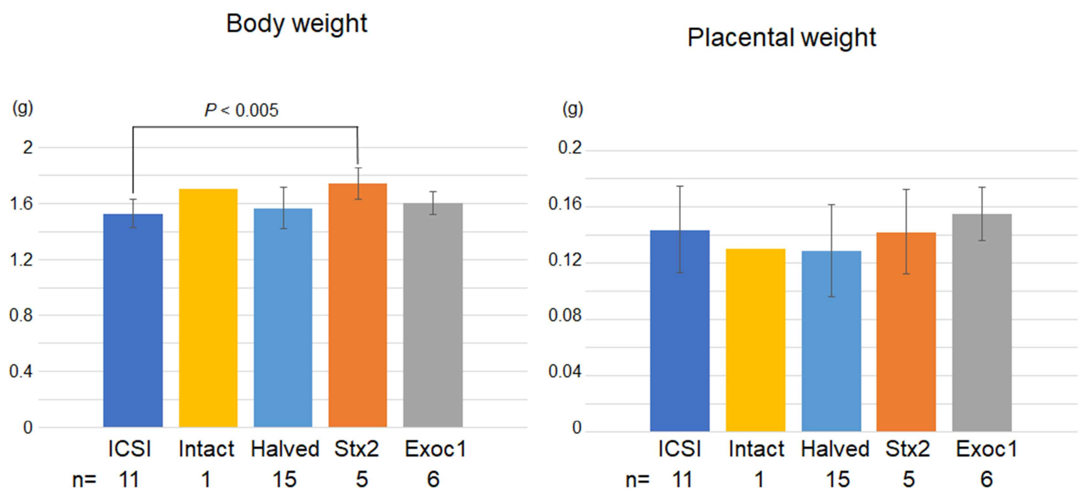


**Figure S1. Body and Placental Weights of Mouse Pups Born Following Spermatocyte Injection** The mean body and placental weights of spermatocyte injection-derived pups were not significantly different from those of ICSI-derived pups, except for the body weight of pups derived from *Stx2*-deficient spermatocytes ($P < 0.005$ by Student's *t*-test). As only one pup was born after experiments using normal size oocytes, this group was omitted from the statistical analysis. Error bars indicate standard deviations. ICSI, intracytoplasmic sperm injection.

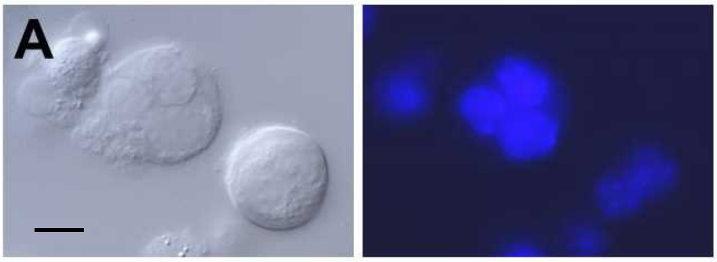


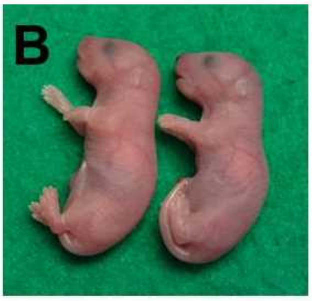


Birth of mice from Exoc1 conditional knockout mice

| No. oocytes cultured | No. (%) 2-cells | No. transferred | No.(%) implanted | No. (%) offspring | With mutation |
|---|---|---|---|---|---|
| 96 | 66 (69) | 46 | 11 (24) | 6 (13) | 3 |

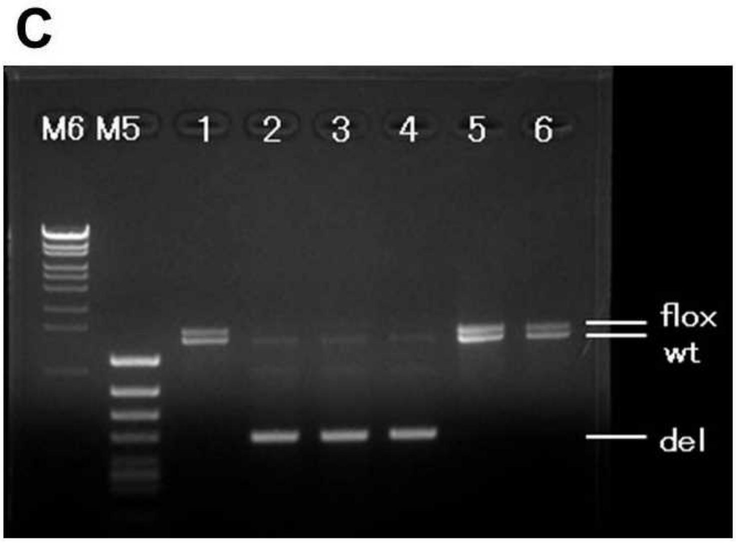


**Figure S2. Birth of mice following injection of oocytes with primary spermatocytes collected from germline-specific *Exoc1*-knockout mice**

(A) Multinucleated cells (syncytial spermatocytes) obtained from an *Exoc1*-knockout male mouse. Each multinucleated cell contained 2–4 spermatocyte nuclei. Differential interference contrast microscopy (left) and Hoechst-staining (right) images. Bar = 10 $\mu$m.

(B) Mice born from *Exoc1*-knockout spermatocytes.

(C) Polymerase chain reaction analysis in mice born from *Exoc1*-knockout spermatocytes. Mice #2, #3, and #4 carried the *Exoc1*-knockout allele (del) while mice #1, #5, and #6 did not. Three pups that did not carry the *Exoc1* mutation were most likely derived from spermatogonia that escaped the Cre-induced *Exoc1* deletion.

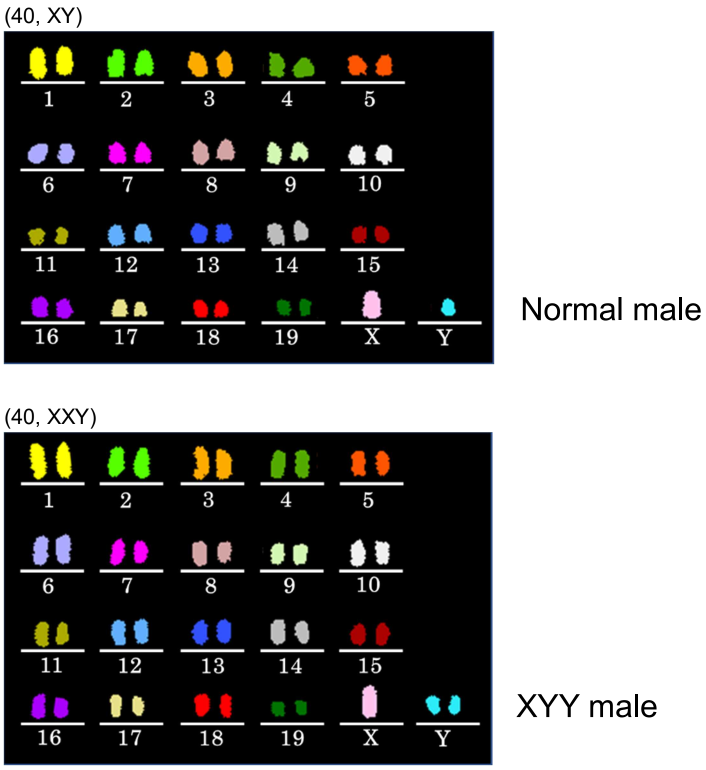


**Figure S3. Chromosomal Multicolor FISH Analysis of the Offspring Derived from Wild-type Spermatocytes**

Of the three males analyzed, one had XYY sex chromosomes. No abnormalities were found in autosomes in these mice.

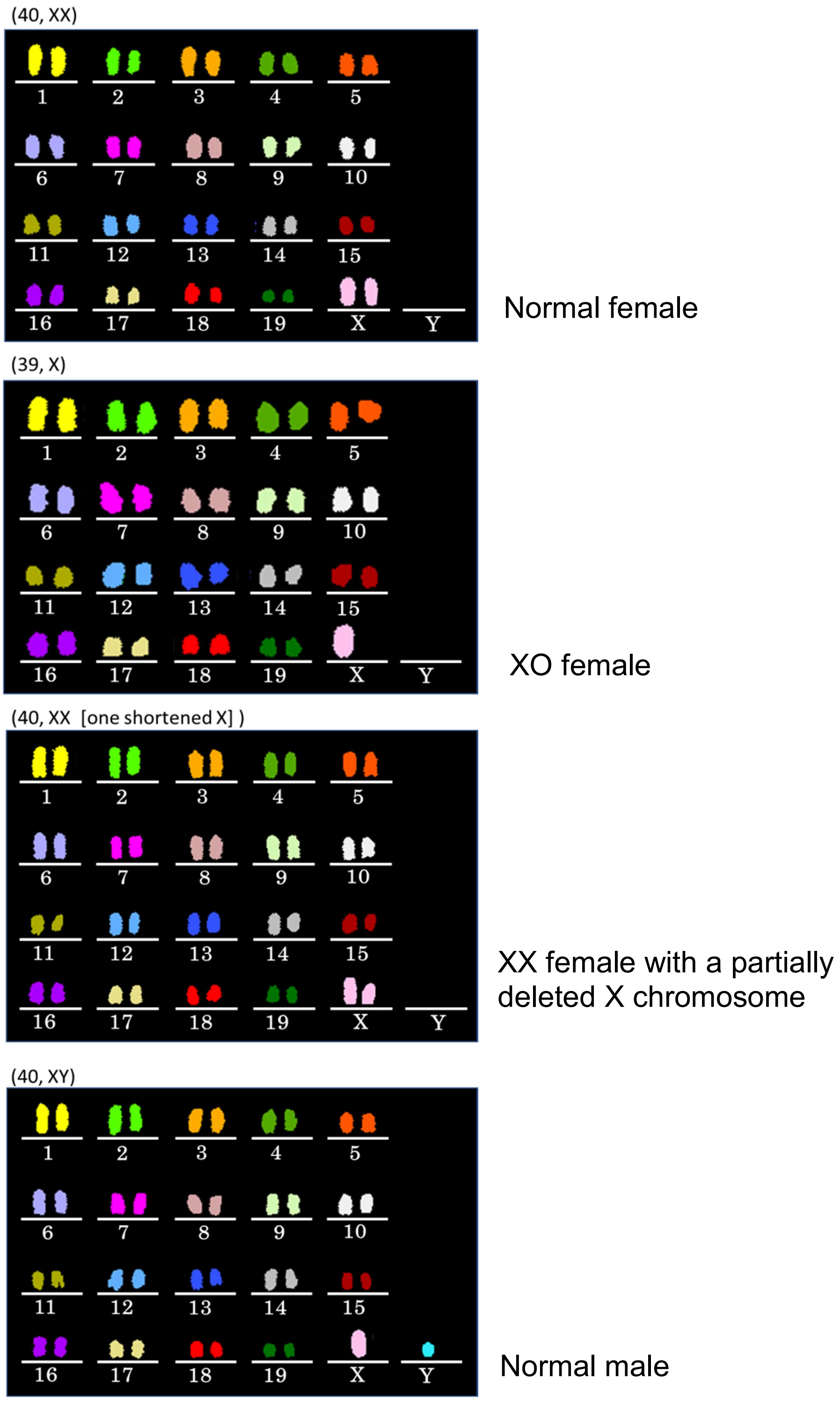


**Figure S4. Chromosomal multicolor FISH analysis of the offspring derived from *Stx2*-deficient spermatocytes**

No abnormalities were found in autosomes, but there were two cases of sex chromosomal abnormalities.

**Table S1. Chromosomal Abnormalities Found in MII Oocytes After Injection with Primary Spermatocytes**

| Spermatocyte injection | Ooplasm reduction (halving) | Oocytes analyzed | Oocytes with normal chr. (%) | Oocytes with abnormal chr. (%) | No. of chromosome aberrations (%) | | | | |
|---|---|---|---|---|---|---|---|---|---|
| | | | | | Premature chromatid separation | Aneuploidy | Condensed chromatin mass | Break | Premature chromosome separation + Break |
| – | – | 39 | 38 (97)[a] | 1 (3) | 1 (3) | 0 (0) | 0 (0) | 0 (0) | 0 (0) |
| + | – | 59 | 1 (2)[a′,b] | 58 (98) | 51 (86) | 3 (5) | 3 (5) | 0 (0) | 1 (2) |
| + | + | 62 | 13 (21)[b′] | 49 (79) | 40 (65) | 6 (10) | 2 (3) | 1 (2) | 0 (0) |

[a vs a′] $P < 0.0001$; [b vs b′] $P < 0.0001$ (Fisher's exact probability test). chr, chromosomes.

**Table S2. Birth of Offspring Following Transfer of Embryos Derived from Injection of Primary Spermatocytes**

| Ooplasm | No. of oocytes cultured | No. (%) 2-cells | No. transferred | No. (%) implanted | No. (%) offspring |
|---|---|---|---|---|---|
| Control | 34 | 25 (74) | 16 | 0 (0) | 0 (0) |
| | 23 | 18 (78) | 18 | 2 (11) | 0 (0) |
| | 33 | 27 (82) | 27 | 4 (15) | 0 (0) |
| | 47 | 35 (74) | 35 | 6 (17) | 1 (3) |
| Total | 137 | 105 (77) | 96 | 12 (13) | 1 (1)[a] |
| Halved | 32 | 22 (69) | 22 | 15 (68) | 8 (36) |
| | 21 | 17 (81) | 9 | 6 (67) | 3 (33) |
| | 40 | 33 (83) | 33 | 13 (39) | 3 (9) |
| | 11 | 11 (100) | 11 | 8 (73) | 2 (18) |
| | 18 | 15 (83) | 15 | 13 (87) | 1 (7) |
| Total | 122 | 98 (80) | 90 | 55 (61) | 17 (19)[a′] |

[a vs a′] $P$ < 0.0001 (Fisher's exact probability test).

**Movie S1. Preparation of a Half-sized Oocyte at the GV Stage**
About half of the ooplasm was removed using a glass pipette. The medium contained cytochalasin D and a high concentration of NaCl to increase the survival rate of treated oocytes.

**Movie S2. Three-dimensional (3D) Reconstruction of Kinetochores and Chromosomal Dynamics**
Chromosomes and centromeres were 3D-reconstructed using major satellite–mClover (centromeres) and H2B–mCherry (chromosomes) marker signals. Magenta and cyan colors indicate the chromosomes from oocytes and spermatocytes, respectively. Red spots indicate the centromeres of univalent-like chromosomes, which underwent predivision at anaphase.

**Movie S3. High-resolution Live Imaging of Kinetochores and Chromosomes**
Oocytes expressing major satellite-mClover (centromeres, green) and H2B-mCherry (chromosomes, red) were used for centromere tracking analysis, shown in **Fig. 2**. Maximum intensity Z-projection images are shown after peak enhancement and background subtraction for centromere signals. Times before anaphase onset are shown in h:min. White arrowheads indicate a univalent-like chromosome that underwent premature segregation of sister chromatids (predivision) at anaphase. Note that in the halved oocyte, one of two anaphase chromosome masses was out of the image stack from times 0:10 to 0:20. Scale bar = 5 μm.

**Movie S4. Injection of Spermatocytes from a *Repro34*-bearing Male Mouse into Half-sized MI Oocytes**
Spermatocyte nuclei were separated from a multinucleated spermatocyte mass. Each nucleus was injected into a half-sized MI oocyte.
